# The mRNA-bound proteome of *Leishmania mexicana*: novel genetic insight into an ancient parasite

**DOI:** 10.1101/592402

**Authors:** Luis M. de Pablos, Tiago R. Ferreira, Adam A. Dowle, Sarah Forrester, Ewan Parry, Katherine Newling, Pegine B. Walrad

**Affiliations:** Centre for Immunology and Infection, Department of Biology, University of York, UK.; Metabolomics and Proteomics Lab, Bioscience Technology Facility, Department of Biology, University of York, UK.; Genomics and Bioinformatics Lab, Bioscience Technology Facility, Department of Biology, University of York, UK.

## Abstract

*Leishmania* parasite infections, termed the leishmaniases, cause significant global infectious disease burden. The lifecycle of the parasite embodies three main stages that require precise coordination of gene regulation to survive environmental shifts between sandfly and mammalian hosts. Constitutive transcription in kinetoplastid parasites means that gene regulation is overwhelmingly reliant on post-transcriptional mechanisms, yet strikingly few *Leishmania trans*-regulators are known. Utilizing optimised crosslinking and deep, quantified mass spectrometry, we present a comprehensive analysis of 1,400 mRNA binding proteins (mRBPs) and whole cell proteomes from the three main *Leishmania* lifecycle stages. Supporting the validity, while the crosslinked RBPome is magnitudes more enriched the protein identities of the crosslinked and non-crosslinked RBPomes were nearly identical. Moreover, multiple candidate RBPs were endogenously tagged and found to associate with discrete mRNA target pools in a stage-specific manner. Results indicate that in *L.mexicana* parasites, mRNA levels are not a strong predictor of the whole cell expression or RNA binding potential of encoded proteins. Evidence includes a low correlation between transcript and corresponding protein expression and stage-specific variation in protein expression versus RNA binding potential. Unsurprisingly, RNA binding protein enrichment correlates strongly with relative replication efficiency of the specific lifecycle stage. Our study is the first to quantitatively define and compare the mRBPome of multiple stages in kinetoplastid parasites. It provides novel, in-depth insight into the *trans*-regulatory mRNA:Protein (mRNP) complexes that drive *Leishmania* parasite lifecycle progression.

## Introduction

*Leishmania* spp. parasites are the causative agent of leishmaniasis, a neglected disease that represents the ninth largest global infectious disease burden [1]. These protozoa have a dixenous lifecycle that transitions between multiple promastigote stages in the sandfly vector to the amastigote stage in the phagolysosomes of mammalian immune cells [2]. Distinct environmental conditions (pH, temperature, nutrient availability) serve as triggers for developmental events. For parasite lifecycle progression, both metacyclogenesis (procyclic to metacyclic promastigote) and amastigogenesis (metacyclic promastigote to amastigote) differentiation processes require tightly coordinated gene regulation [2]. To date, many *cis*- elements but strikingly few *trans*-regulators have been implicated in *Leishmania* developmental progression [3-11]. The identification of *Leishmania trans*-regulators that bind mRNAs in a stage-specific manner lends vital insight into the cellular processes which promote and enable parasite survival.

Regulation of gene expression requires fine-tuned, coordinated mechanisms that respond to shifting environmental conditions. Kinetoplastid parasites rely almost exclusively upon post-transcriptional gene regulatory mechanisms due to their constitutive transcription of Pol II-driven polycistronic gene arrays [2]. Accordingly, RNA binding proteins (RBPs) are overrepresented in the proteome of these organisms in line with their role as the primary gene regulators. Such *trans*-regulatory RBPs dynamically bind to mRNA forming ribonucleoprotein complexes (mRNPs) that regulate the trafficking and processing of mRNA molecules from synthesis to decay [12]. Environmental pressures stimulate swift changes in mRNP localization, composition and function that accelerate rates of mRNA translation, decay, or sequestration to intracellular granules in response [13].

Previous large-scale isolations of mRNA-bound proteomes have identified candidate regulators in yeast, flies, mice, and humans [14,15]. More recently, kinetoplastid parasite mRNP investigations yielded 155 RBPs in *Trypanosoma brucei* monomorphic bloodstream forms and 128 *Leishmania donovani* RBPs in axenic amastigote forms [16,17]. These studies were refined in scope to only one lifecycle stage, yet importantly confirm that mRNA-bound factors in kinetoplastid cells include proteins without canonical RNA-binding motifs [14-18].

Here we present the mRNA-bound as well as whole cell proteomes isolated from *L.mexicana* procyclic promastigote (‘PCF’), metacyclic promastigote (‘META’) and amastigote (‘AMA’) stage parasites. We have identified over 1,400 mRNA bound proteins (RBPs) represented by at least two unique peptides in both UV-crosslinked (XL) and non-crosslinked (nonXL) samples, the XL samples being magnitudes higher in overall enrichment due to the covalent bonds strengthening interactions. The isolated mRNA-bound proteomes are differentially enriched and over 250 RBPs exhibit stage-specific expression. In addition, the whole cell proteomes of these stages were also quantitatively identified in triplicate for comparative use in this study and represent an essential resource for the *Leishmania* and eukaryotic research community. Of 8,144 predicted proteins, our analysis identified over 2,400 with at least 2 high quality unique peptides, of which nearly half fluctuate in expression levels throughout the lifecycle. Of interest, bioinformatics and biochemical analyses indicate only a minority of identified proteins display expression patterns similar to their encoding transcripts. Furthermore, the expression of an mRNA binding protein does not correlate to RNA association. Importantly, these findings may implicate post-translational modifications in the fluctuations of RNA binding potential and relative stability of candidate *trans*-regulators as well as overall cellular translational capacity and activity.

## Results

### Isolation and validation of *L.mexicana* life cycle stages

For the study of the different mRNA binding proteomes (mRBPomes) during the *L.mexicana* lifecycle, 4 biological samples corresponding to the 3 main lifecycle stages were isolated and molecularly verified. These were low passage (< P3), M199 media culture-derived procyclic promastigotes (‘PCF’), Grace’s media culture-derived metacyclic promastigotes (‘META’), 24 hour post-infection (24hr pi) macrophage-derived amastigotes (‘AMA(MØ)’) and *in vivo* lesion-derived amastigotes (‘AMA(LD)’; Fig 1A). Parasite cells were validated for specific lifecycle stages using distinct molecular markers and biologically-distinguishing features including cell cycle replicative status, resistance to human serum lysis and marker gene expression (Fig 1B-D). The cell cycle analysis of PCF cell cultures mid-log at 5×10^6^ cells/ml showed high replication efficiency (S: 19%, G2/M: 35%), sensitivity to human serum incubation and heightened *Histone h4* (*h4*) transcript expression (Fig 1B, C and D). We introduce *Histone H4* (*LmxM.36.0020*) here as a novel transcript marker of PCF stage *L.mexicana* parasites, investigated due to its specific expression in sandfly-derived *L.major* PCF cells [19]. Stationary META form parasites were harvested at ~4×10^7^ cells/ml 7 days post-incubation in Grace’s media as described [20] and validated by reduced replication and protein synthesis [21](S: 3%; G2/M: 19%), resistance to human complement lysis [22] and heightened expression of *sherp* transcript [23](Fig 1B, C and D). Intracellular amastigote-stage parasites were isolated 24hr post-infection (24 h.p.i.) of cultured J774.2 macrophages, AMA(MØ), or from mouse rump lesions 4 months post-inoculation [24], AMA(LD), with serum-resistant META cells and gradient-purified as described (Experimental Procedures) [6]. The 24 h.p.i. timing was selected as RNA levels are remodeled and differentiation into AMA(MØ) forms in the lysosomal environment is nearly complete [25]. Purified amastigote-stage parasites were validated by electron microscopy (data not shown), heightened *amastin* (*LmxM.08.0840*) mRNA expression and cell cycle analysis that showed increased replication efficiency relative to META cells (Fig 1B and D). Remarkably, FACS analysis of AMA(MØ) and AMA(LD) showed near-identical cell cycle profiles, intermediate between PCF and META replication efficiencies (Fig 1B).

**Figure 1.**
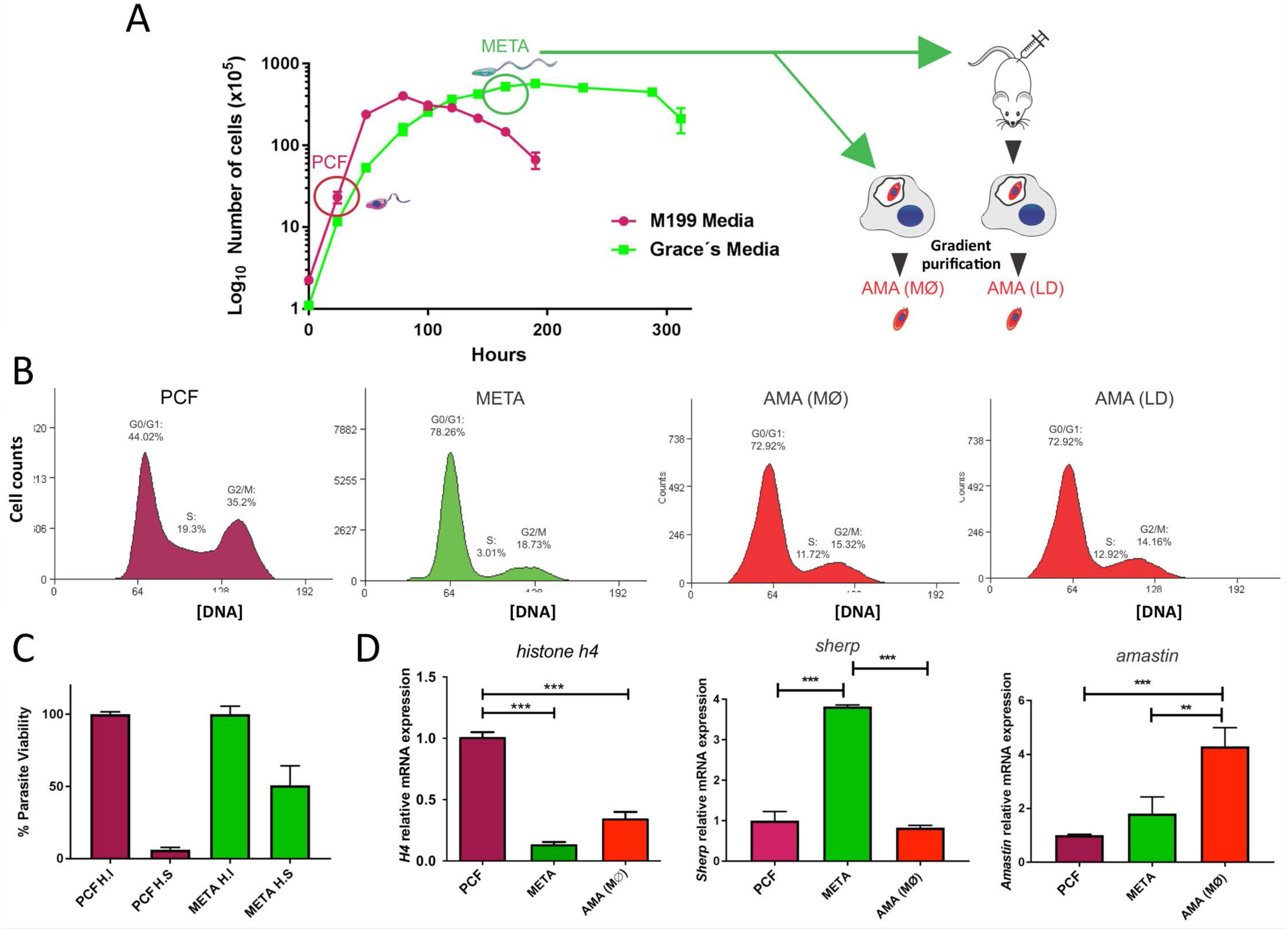
Isolation and validation of *L.mexicana* lifecycle stages. (A) Cumulative growth curve of *L.mexicana* in M199 (purple) and Grace’s media (green). Harvested procyclic (PCF, purple circle) and metacyclic-enriched promastigote cultures (META, green circle) were tested for stage-specific traits. Metacyclic-enriched cultures were used to infect immortalized macrophages (J774.2 MØs) and mice to generate AMA (MØ) and AMA (LD) amastigote forms, respectively. Amastigotes forms (24hr post-infection (pi) for AMA (MØ) and 4 months pi for AMA(LD)) were gradient purified. (B) FACS analyses compare DNA profiles of PCF, META, AMA (MØ) and AMA (LD) cells. Cell cycle profiles demonstrate PCF are highly proliferative, META are relatively quiescent and distinct AMA populations display a near-identical intermediate replicative capacity. (C) Human serum resistance expressed as percentage parasite viability (y-axis) of PCF (purple) and META (green) parasites treated with Human Serum (H.S) or Heat Inactivated human serum (H.I). (D) Validation of each isolated *L.mexicana* stage by heightened mRNA expression of *histone h4*, *sherp* and *amastin* relative to *nmt* transcript levels presented as mean ± SE from 3 experimental replicates. One-way ANOVA and Tukey’s post-test were conducted; **p<0.01, ***p<0.001.

### *In vivo* Capture of *L.mexicana* RBPs

Fig 2A illustrates the procedure used to isolate the mRBPomes. Indeed, up to 91% of cells showed wildtype morphology post-UV crosslinking (XL) with a small portion of non-crosslinked (nonXL) parasites able to restore culture growth after 120s irradiation (Fig 2B). Thus, this length of UV-exposure was selected for the mRNA-interactome capture experiments, with the mRNA subsequently isolated as previously described [18]. All experimental conditions were optimized for large-scale harvests in order to isolate enough mRNA bound proteins (RBPs) for reliable, quantitative mass spectroscopy for each replicate. To round these analyses, relative enrichment of the mRBPomes were compared to whole cell [26] proteomes from each lifecycle stage; PCF, META, AMA(MØ). All peptide pools were evaluated simultaneously in triplicate on the Orbitrap Fusion™ mass spectrometer to quantitatively determine relative intensities of peptides in the mRBPomes and WC proteomes. The purity of the oligo(dT)-derived XL and nonXL mRNA was evaluated by relative elution of *18S* ribosomal RNA compared to the constitutively-expressed *N-myristoyltransferase* (*nmt; LmxM.31.0080*) mRNA as reference for overall mRNA recovery (Fig 2C) [27]. The results show a negligible amplification of *18S* rRNA relative to input and a substantial fraction of the *nmt* mRNA recovered after poly-A RNA isolation.

**Figure 2.**
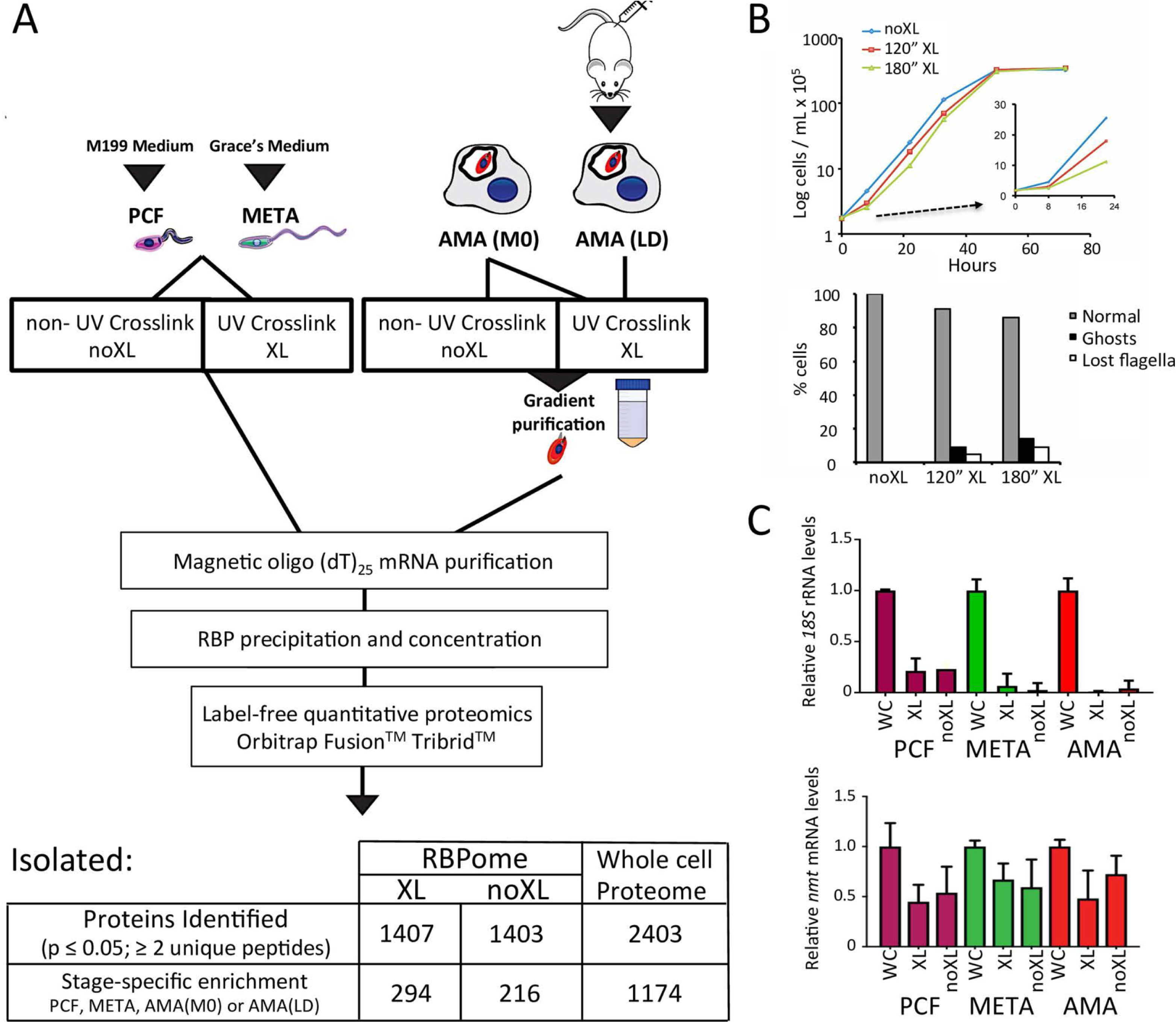
Overview of the mRNA-interactome capture workflow. (A) *Leishmania* cells from verified lifecycle stages are UV-irradiated *in vivo* creating covalent bonds between mRNA and bound proteins in crosslinked (XL) versus non-crosslinked cells (nonXL). Cells are then purified, lysed and mRNP complexes are isolated using oligo(dT)-labeled magnetic beads. Isolated RBPs were precipitated and processed for label-free, quantified mass spectrometry. Numbers indicate identified proteins filtered for triplicate-consistent, high quality reads with at least 2 unique peptides. (B) Growth curve and cell morphology after 120” or 180” UV-irradiation with optimised equipment. (C) Relative RNA recovery of *18S* and *nmt* transcripts after oligo(dT) magnetic bead mRNA purification in XL, nonXL and WC lysates in PCF (purple), META (green) and AMA (MØ; red) samples.

### Whole cell and RNA-bound proteomes of the main *Leishmania* lifecycle stages

The captured RBP peptides were analysed by high-resolution mass spectrometry with a filter minimum of 2 unique peptides identified per protein and quantified using peptide precursor ion intensities (p ≤ 0.05). A total of 1,407 RBPs were identified in the XL samples (PCF, META, AMA(MØ), AMA(LD)) with 294 proteins significantly enriched at distinct stages, while 1,403 RBPs were isolated from the nonXL peptide samples (PCF, META, AMA(MØ)) with 216 proteins displaying stage-specific association (Fig 2A). This number is consistent with current RBP numbers in other eukaryotic systems [15]. The RBPomes of 4 distinct *Leishmania* lifecycle stages have been pooled; equivalent to 4 distinct mammalian cell types, and our experimental method removes potential competition and quantitative limitations introduced by ion labeling (Experimental Procedures). In addition, mass spectrometry of whole cell [26] *L.mexicana* proteomes of each stage (PCF, META, AMA(MØ)) yielded 2,403 proteins identified, 1,174 of which are significantly enriched at one lifecycle stage (Fig 2A). These data are available in Supplementary S1 Table. Confirming the integrity of our proteomes, principal component analysis (PCA) using relative protein quantification derived from unique peptide intensities was used to examine relatedness between the different biological samples and the variability among replicates (Fig 3A). Notably, the triplicate proteomic samples cluster within discrete lifecycle stages for XL, nonXL and WC proteomes displaying low variance among replicates and reproducibility of the results (Fig 3A). These data infer reliable and distinct protein enrichment and identities for each lifecycle stage. In accordance with the FACS analyses of Fig 1B, the PCA values for the AMA(LD) XL mRNA binding proteome is most similar to that of the AMA(MØ) XL, validating the use of AMA(MØ) for subsequent mRNP analyses. When combined, the WC proteomes cluster separately compared to the XL and nonXL mRBPomes, indicating that the isolated RBPs represent an enriched fraction with distinct relative intensities from that of the whole cell proteomes (Supplementary S1A Fig). Supporting the validity of the mRBPome, PCA analyses indicate the total proteomes, regardless of lifecycle stage, are more similar to each other in peptide identities and intensities than to any of the mRNA-selected proteomes (mRBPomes).

**Figure 3.**
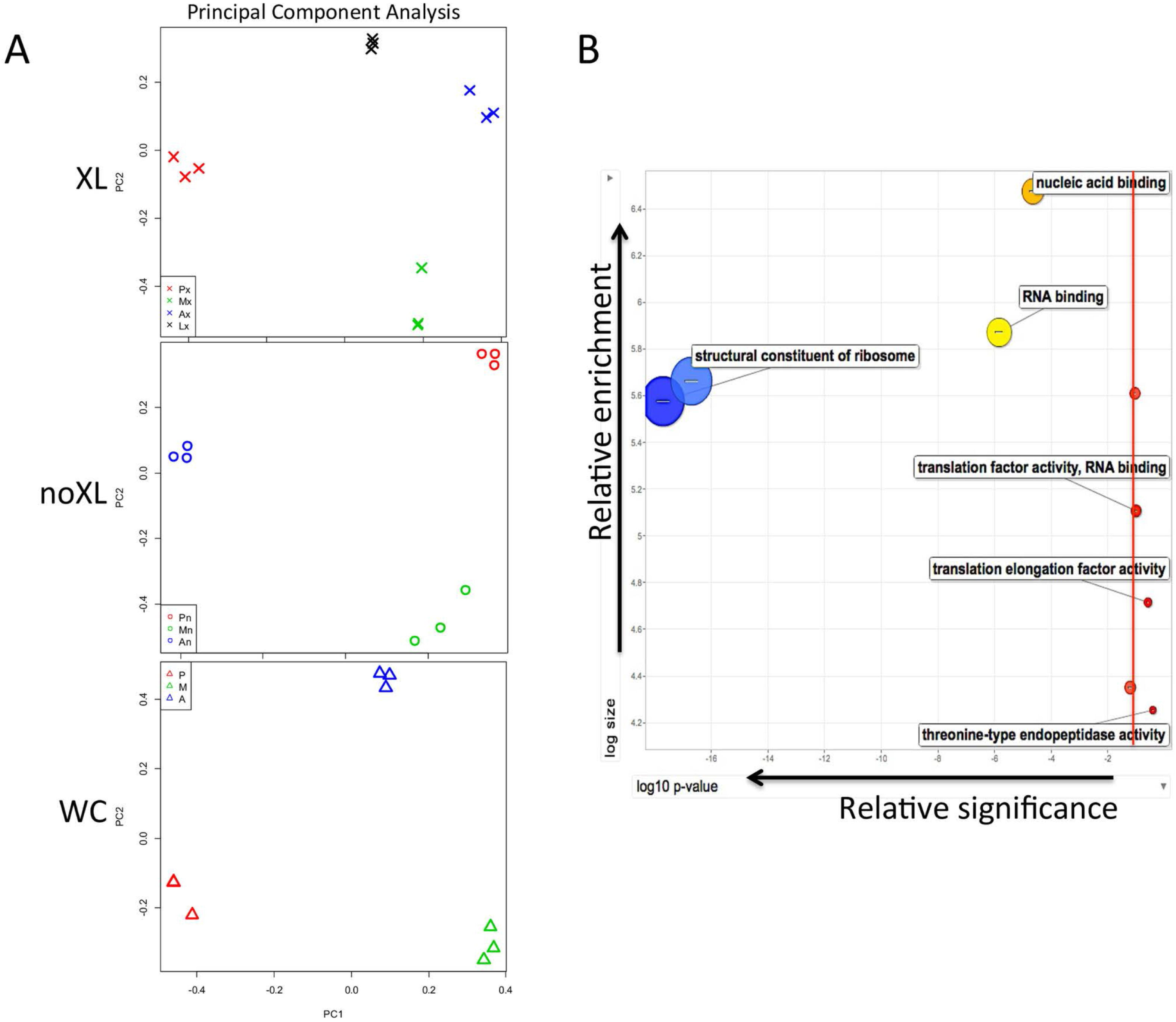
Variability and GO term enrichment among proteomes. (A) Principal component analysis (PCA) using relative protein quantification derived from unique peptide intensities was used to examine relatedness between the different biological samples and the variability among replicates. Notably, the triplicate proteomic samples cluster within discrete lifecycle stages for XL, nonXL and WC proteomes displaying low variance among replicates and reproducibility of the results. Colors indicate lifecycle stage of samples: PCF (red), META (green), AMA(MØ) (blue) and AMA(LD) (black). XL Samples (X) are Px = PCF XL, Mx = META XL, Ax = AMA(MØ) XL and Lx = AMA (LD) XL. NonXL Samples (O) are Pn = PCF nonXL, Mn = META nonXL and An = AMA(MØ) nonXL. WC Samples (Δ) are P = PCF WC, M = META WC and A = AMA(MØ) WC. (B) Within the specific 234 RBPs Gene Ontology (GO) term enrichment analyses again indicated that this specific *L.mexicana* subset was enriched in the following terms by the following order: structural constituent of the ribosome (p = 4.67e^−10^), RNA-binding (p = 1.25e^−6^), nucleic acid binding (p = 7.2e^−4^) indicating that the Molecular Function subfraction specific to the *L.mexicana* RBPome is overwhelmingly enriched in RNA-related terms. Red line indicates p ≤ 0.05.

The specificity of our RNA binding proteomes (XL, nonXL) relative to the whole cell [26] proteome are further confirmed by Gene Ontology (GO) term enrichment analyses which demonstrate that significantly-enriched terms (p ≤ 0.05) from the Molecular Function subset were reliably RNA-related; restricted to ribosomal components, RNA-binding and nucleic acid-binding with translation-related factors ranking somewhat less significantly (Fig 3B). Despite a somewhat abridged proportion of annotation for the *L.mexicana* proteome, with 519 of the 1410 RBPs being hypothetical proteins, known protein identities within the isolated total mRBPome indicate appropriate enrichment.

Of interest, the number and identities of the RBPs within the XL versus nonXL mRBPome samples are remarkably similar, yet the relative peptide intensities are differentially-enriched (Supplementary S2 Fig). This differential enrichment is due to the strength of RBP assocation in the XL samples being based upon the number of covalent bonds formed when direct amino acid:mRNA interactions are irradiated (254nm) while the nonXL protein:mRNA associations are reliant upon the relative affinity of each RBP for mRNA [28]. Accordingly, while the overall XL proteome is greatly enriched (≥ 10 fold) relative to the nonXL samples, the nonXL RBPome is enriched for RNA helicases which exhibit particularly strong binding in clamped conformation [29]. The fact the protein identities are overwhelmingly conserved between the XL and nonXL RBPomes supports the biological relevance of our results and deviates from previous kinetoplastid studies, which have used the nonXL samples as negative controls [16]. Refined UV-crosslinking combined with the high sensitivity of the Orbitrap Fusion™ mass spectrometry system enables a deeper, quantified identification of nonXL without competing with the more abundant XL RBPome. Importantly, running each sample independently, label-free and concurrently prevented peptide competition for ion labeling and signal quenching, enabling quantitative comparison despite the significantly higher overall intensities of the XL versus the nonXL mRBPome samples. We further verified the stringency of all mRNA harvests (WC, XL and nonXL) through the lack of contaminating rRNA levels (Fig 1C).

### Proteomic comparisons reveal stage-specific distinctions

Filtered WC, nonXL and XL proteomes of each stage were compared using log_2_ fold change of peptide ion intensities ≥ 10^6^ among biological samples to isolate factors which are stage-specifically enriched (Supplementary S2 Fig). Importantly, filtering by intensity enables visualization of distinct protein enrichment but potentially excludes functionally-relevant proteins below this threshold. Comparison of the WC proteomes yielded 69, 50 and 27 proteins which are specifically enriched in PCF, META or AMA(MØ) stage parasites, respectively, with the majority common to all lifecycle stages. In contrast, the majority of mRNA binding proteins are not common to all lifecycle stages but are differentially enriched and stage-regulated in both the nonXL and XL RBPomes (Supplementary S2A Fig). A complementary way of examining this data is provided in Supplementary S2B Fig; which compares the same protein enrichment analyses of the WC, XL and nonXL proteomes of each lifecycle stage (Supplementary S2B Fig).

Comparison between the predicted RBD-containing proteome of the *L.mexicana* genome (tritrypdb.org) and the isolated RBPome (XL) highlights some interesting distinctions (Fig 4A). A significant reduction of zinc-finger domain proteins is evident in the isolated RBPome; with a 50% decrease in CCCH domain protein enrichment from 8% to 4%. This is consistent with the relative ‘silence’ of zinc finger proteins in mass spectroscopy. Supporting the specificity of our data, the mRBPomes isolated from the highly replicative PCF stage displayed a large increase in the number of RBPs homologous to basal translational machinery. Overall, only 31% of the predicted proteins with RBDs in the whole genome are detected in our RBPome. The most likely explanation for this are the multiple timepoints within the *L.mexicana* lifecycle and *in vivo* conditions beyond the scope of this study.

**Figure 4.**
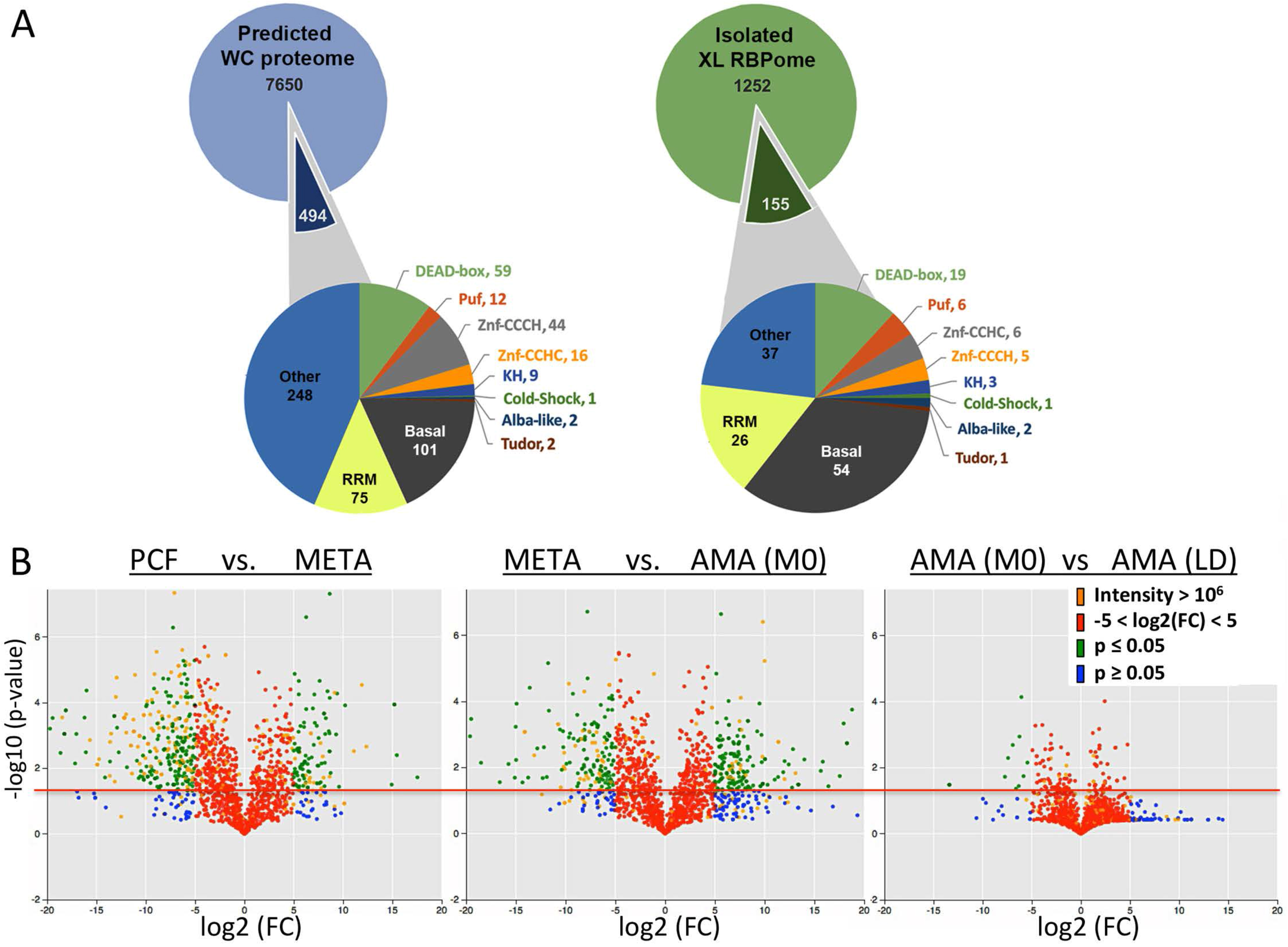
*L.mexicana* lifecycle whole cell and RBPome comparisons. (A) The number of proteins containing common RNA-binding domains (RBDs) in the predicted *L.mexicana* proteome (Tritrypdb.org) and XL-RBPome are presented. 494 out of 8144 genes in the total proteome and 155 out of 1407 proteins in the XL-RBPome contain known RBDs. Domain classes labeled as ‘Other’ represent proteins with either additional, less abundant RBDs not listed separately in the diagram or weak conservation of known RBDs and ‘Basal’ proteins have homology to the translational and splicing machinery. The RBPome RBD total is higher than the number of RBPs identified due to the presence of multiple domain classes in single proteins. (B) Volcano plots were generated in Python using p-values derived by conducting independent t-tests (3 replicates in stage 1 vs 3 replicates in stage 2). No multiple testing was conducted as the plots are for data visualization. The log_2_(FC) is log_2_((mean exp. condition 2 +1)/(mean expression condition 1 +1)). Red line indicates p ≤ 0.05; protein identities above are significantly enriched in one lifecycle stage. Blue circles indicate identities below this threshold, green above. Red circles indicate proteins in the range of −5 < log_2_(FC) < 5 and orange circles indicate highly expressed factors with intensities ≥ 10^6^.

To more closely compare the mRNA-bound proteomes between lifecycle stages without the exclusion of RBPs with lower intensities, XL proteomes of each stage were examined using log_2_ fold change of peptide ion intensities (Fig 4B). Volcano plots represent the change in abundance (x-axis) versus significance (y-axis) of individual RBPs captured among the XL biological samples (Fig 4B). The proteins of intensity ≥ 10^6^ that are analyzed in Fig 4A can be observed here in orange versus the remaining identities. What is evident is that these proteins of intensity ≥ 10^6^ make up the minority of protein identities overall, with identities above the red line displaying significantly distinct association with mRNA between the compared lifecycle stages. Identities in green have reduced intensities (≤ 10^6^) but larger stage-specific shifts in protein bound to mRNA (log_2_(FC) < −5 or > 5). Consistent with the high division rate characteristic of the PCF stage (Fig 1B), the PCF-specific XL RBPome is enriched for factors implicit in translation relative to the quiescent META stage XL mRBPome. Interestingly, large distinctions in relative enrichment are observed between the XL RBPomes enriched in META versus AMA(MØ), which are temporally separated by only 24 hours. These RBPs are likely implicit in the differentiation potential [30] and initiation AMA(MØ) of amastigogenesis. Remarkably, the AMA stage samples display the least distinction of the XL mRBPomes despite the different host environments of an immortalized macrophage cell line versus an *in vivo* lesion 4 months post-infection (Fig 4B). This finding contrasts the Venn Diagram comparison of the highly enriched AMA(MØ) and AMA(LD) RBPomes (Fig 4A) and suggests that while the identities of a minority of highly enriched factors (intensities > 10^6^) are distinct, the overall RBPomes of AMA(MØ) and AMA(LD) are remarkably similar. Combined with the near-identical cell cycle profile (Fig 1B), the volcano plot profiles of AMA(MØ) and AMA(LD) (Fig 4B) support cultured macrophages (MØs) as a useful model to investigate potential *trans*-regulators of *L.mexicana* differentiation *in vivo*.

### mRNA and protein expression versus RNA-binding activity of RBPs

To validate proteomic results biochemically, specific RBPs of interest were endogenously HA-tagged on the N-terminus to examine expression dynamics as controlled by endogenous 3’UTRs. Western blots confirmed stage-specific protein expression that corresponds with mass spectrometry results for RBP16 and UBP1 (Supplementary S3 Fig). In comparison with mRNA expression, protein expression of these RBPs displayed interesting variances. While DRBD2 shows a relatively close correlation between transcript and protein levels, levels of SUB2, RBP16, DRBD3 and UBP1 proteins do not correspond well to encoding transcript levels (Supplementary S3 Fig).

This led to a comprehensive examination of the previously observed phenomenon that *Leishmania* spp. protein and mRNA levels do not closely correspond [7]. Given the scale of our proteomic results, we were able to use a GLM (Generalized Linear Model) of loess algorithm fit with a sliding window approach to visualize potential correlation between our WC (Purple) and XL (Green) proteomic data relative to published transcriptomic data [25] in both AMA(MØ) and PCF lifecycle stages (Fig 5A). This comparison was limited to two lifecycle stages due to the transcriptome data available [25], however the low overall Pearson’s correlation values indicate that neither the whole cell (R^2^ = 0.14) nor RNA-bound (R^2^ = 0.0057) proteome intensities correspond to levels of transcript expression. While a stronger correlation between transcriptomic and whole cell proteomic data is evident in PCF stage cells which are more translationally active than AMA (Fig 1B), there is negligible connection between the transcript expression of an RNA bound protein and its subsequent association with RNA (Fig 5A). These results indicate that in *L.mexicana* parasites, RNA levels are not a strong predictor of whole cell expression or the RNA binding potential of proteins.

**Figure 5.**
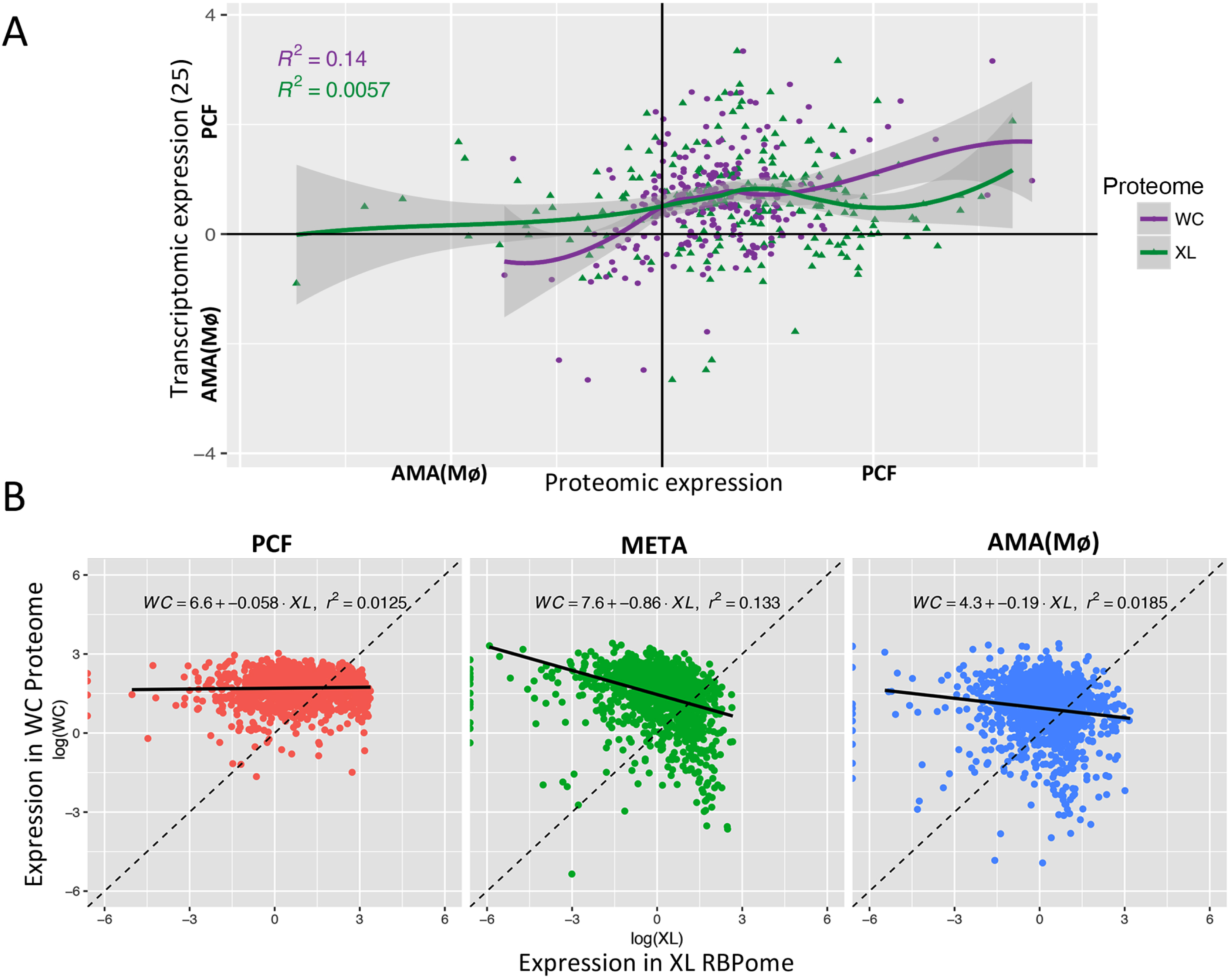
Comparing *L.mexicana* WC and XL Proteomes to Transcriptome data. (A) GLM (Generalized Linear Model) of loess algorithm fit displaying a sliding window approach to visualize relative correlation between the WC (Purple) and XL (Green) proteomic data relative to published *L.mexicana* transcriptomic data [25] in AMA(MØ) vs PCF lifecycle stages. Grey shading = S.E. Relevant statistics provided in Table 1. (B) A linear regression fit against all proteins which meet XL/WC ratio, Lm [WC~XL]. Expressed proteins are more likely to bind mRNA in PCF than META with AMA intermediary. Likelihood for expressed proteins to bind RNA correlates well to both replicative and translational efficiency of each lifecycle stage.

This comparison stimulated the question of how well WC RBP expression correlates to RNA binding potential (XL). Fig 5B illustrates a linear regression fit (Lm [WC~XL]) to examine the RNA binding potential (XL RBPome; X-axis) of proteins quantified in our whole cell protein expression (WC RBPome; y-axis) which meet XL/WC ratio Lm [WC~XL]. As our methods do not exclude the isolation of the translational machinery in our investigation, the high expression levels of translation factors influence this proteomic analysis. While stages with high replication rates indicate a stronger correlation between protein expression [26] and RNA binding (XL), overall the correlations are lower than expected (Fig 5B). Data from Figs 1B and 4B also display influences relevant to translational activity as AMA cells are intermediary between PCF and META in both replication and translational efficiency with little correlation between RNA binding protein expression [26] and RNA association (XL) in the translationally-repressed, quiescent META stage parasites (Fig 5B). These data indicate that expression of an RNA binding protein [26] is not a strong indicator of its RNA binding (XL). Importantly, this implies another level of regulation modulates or alters the RNA binding potential of RBPs. Given the strong evidence of stage-regulated post-translational modifications in this system [31], PTMs likely contribute to RNA binding potential.

### Validation of novel RNA binding proteins in *L.mexicana*

In order to functionally validate our mRBPome, multiple RBP candidates previously-uncharacterized in *Leishmania mexicana* were endogenously tagged (Fig 6A), RNA immunoprecipitated (RIP) and associating transcripts were sequenced (data not shown). Top putative target transcripts for each RBP were validated and examined for stage-specific association by additional RIPs (Fig 6A) and subsequent qRTPCRs of whole cell (Fig 6B) versus RBP-associated transcript levels (Fig 6C) in both PCF and META stage parasites. As expected, candidate RBPs associate with specific pools of transcript targets and this association can be stage-regulated. Of interest, some transcript targets are shared between RBPs and stage-specific fluctuations in mRNA-affinity can be target-specific. Remarkably, one of the more interesting RBP candidates, GAPDH protein, is expressed at relatively constant levels between PCF and META stage parasites, yet selects distinct transcript target pools in these stages. This may suggest that the availability of a given RBP to bind mRNA can be altered in a bespoke manner to adjust its specificity, rather than a simple fluctuation in RNA binding capacity, in a stage-specific manner. Alternatively, and not exclusively, it may be that the structure and exposure of regulatory elements within mRNA targets change during the lifecycle. On the whole, this molecular data supports and extends our bioinformatic findings that the whole cell expression of an RNA binding protein in a given lifecycle stage does not guarantee its association with target transcripts. Instead, RBP expression merely establishes the potential to bind and regulate mRNA, which is then subject to cellular context.

**Figure 6.**
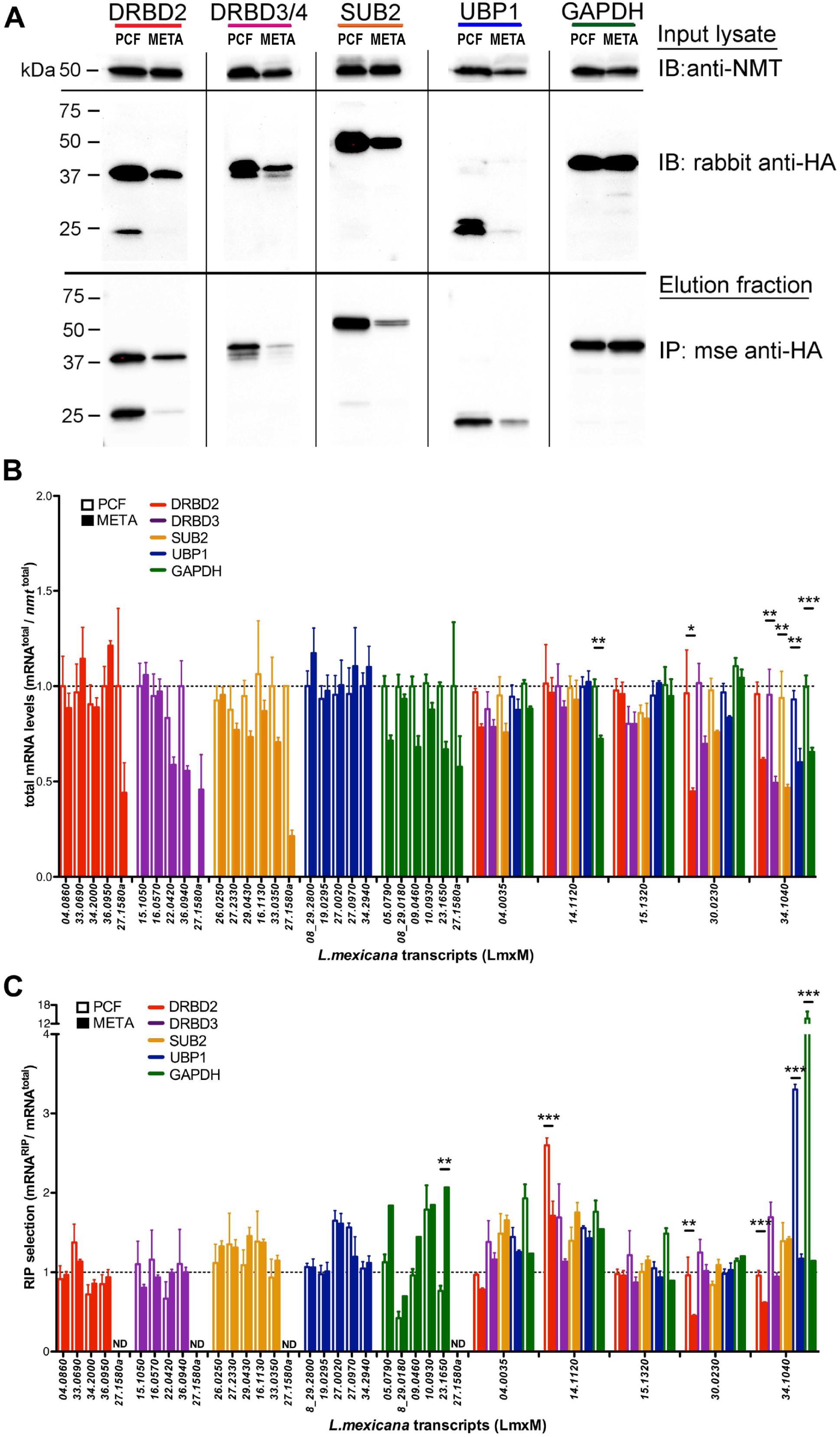
Validation of novel *L.mexicana* RBPs. A. Western blots (anti-HA) of whole cell lysate (Input) and immunoprecipitations (Elution) of endogenously-tagged RBP candidates show specific isolation in both PCF and META stages. B. Whole cell expression of candidate transcript targets are shown in both PCF and META stages, normalized to *nmt* transcript expression. C. HA-RBP RIP-selected mRNA levels are shown relative to WC levels (B) for each target RNA. All qRTPCR data are presented as mean ± SE from 3 biological replicates. Two-way ANOVA test and Bonferroni between log and stationary phase samples indicate a stage-regulated association with target transcripts for some RBPs; \**p*<0.05, \*\**p*<0.01, \*\*\**p*<0.001. ND = Not Detected.

## Discussion

Gene regulation in *Leishmania* spp. parasites is overwhelmingly post-transcriptional as genes are constitutively transcribed in polycistronic tandem arrays [2]. Despite a strong emphasis upon *trans-*regulators as the primary gene regulators, remarkably few have been characterized or validated in these parasites. Here we present a comprehensive, quantitative list of the mRNA-bound proteomes as well as the whole cell proteomes of the 3 main lifecycle stages; procyclic promastigotes, metacyclic promastigotes and amastigote stage *L.mexicana*. We isolated and molecularly validated distinct stages via published criteria including growth in bespoke media, replication efficiency, relative resistance to human serum complement and heightened expression of stage-specific gene markers [20-23]. As a result, we present a comprehensive, updated method for obtaining large quantities of validated, distinct *L.mexicana* lifecycle stages that provide useful insight for the research community. The proteomes of different stages display remarkable protein expression diversity in line with the different environments they inhabit. Cell-type specific gene expression is particularly common in parasites as a protein that is key for survival in one lifecycle stage may represent a vulnerability in the next. We present here the foremost comprehensive RNA binding (XL, nonXL) and whole cell [26] proteomes as yet available in *Leishmania* parasites as a significant resource for the research community.

Several publications have previously described lifecycle-specific proteomes in *Leishmania* spp. parasites [6,7]. Notably, Paape et al. previously published proteomes derived from logarithmic promastigote and macrophage-derived amastigote stage *L.mexicana* in search of potential virulence factors [6]. This study identified enriched octomers within 3’*UTR*s that may contribute to stage-specific protein expression. Further to this, a direct comparison by Lahav et al. observed that transcriptomes of axenic *L.donovani* promastigotes and amastigotes did not correspond to proteomic data, except during the first hour of axenic amastigogenesis [7]. Our data (Fig 5A) supports this finding albeit on a broader timescale than an hourly time-course and in a different *Leishmania* species, suggesting this lack of correlation between mRNA and protein expression may be common to all *Leishmania*. Certainly transcript expression potentiates *Leishmania* protein expression and thereby provides useful insight into gene expression that would be prohibitively expensive and experimentally challenging to obtain on the proteomic level currently [19].

Recently, there have been two RBPome studies in kinetoplastid parasites which isolated 128 RNA bound proteins from axenically-derived amastigote stage (AXA) *L.donovani* [17], and 155 RBPs from *T.brucei* monomorphic slender bloodstream form (BSF) parasites [16]. These studies each examine a single, mammalian-infective form of the parasite. Direct comparisons between the RBPomes have caveats, given the large divergence between intracellular (*Leishmania*) versus extracellular (*T.brucei*) cells and different isolation methods involved. As the PCF, AXA and BSFs are all proliferative stages, translation factors implicit in cell replication are common to all these RBPomes. Our data examines RBPome association dynamics through lifecycle progression utilising the foremost mass spectroscopy technology. This strategy revealed stage-specific modulation of RBP:mRNA associations indicative of a highly bespoke transcript target selection that is independent of expression levels. Indeed, this complicates the traditional RBP:transcript target paradigm in a meaningful, interesting manner with similarities to transcriptional context.

The scale and filters we have employed for our investigation select the most abundant proteins bound to mRNA. Accordingly, in addition to specific *trans*-acting gene regulators we have isolated the translational machinery of each lifecycle stage. Therefore, the relative replication and translational efficiency of each lifecycle stage has strongly influenced many of our RBPome comparisons; including the observed correlation between proteomic expression and mRNA association. While it is beyond the scope of this study, our data provides significant direction toward *Leishmania* ribosomal analyses. Indeed stage-dependent distinctions in the translational machinery may impact general mechanisms, however the primary objective of these analyses is to identify mRNA-bound factors that regulate gene expression and promote differentiation to human-infective forms. That is, *trans*-regulatory RBPs which coordinate both developmental and virulence-promoting regulons.

Given the depth and sensitivity of our proteomics, we cannot rule out the possibility that some of our factors may be indirectly bound by tight protein:protein interactions with directly-bound RBPs. This may explain the presence of non-canonical RNA associating factors; the validity of which is supported by their consistent isolation in RBPomes from other organisms and the knowledge that the current list of RNA binding domains is not exhaustive [14,16,17,32]. The consistency and quality of our methods and results are evident in the near-match identities of our proteomes isolated from mRNA derived from both *in vivo* crosslinked and non-crosslinked cells of all lifecycle stages, negating artifactual RNA binding as a result of UV-crosslinking. Our analyses compare relative enrichment of factors within the Whole Cell versus mRNA-bound proteomes and found highly abundant proteins which also bind mRNA appear to be limited to ribosomal factors. Thus we conclude that non-specific background proteins were not isolated in our screen.

The data from Fig 4B demonstrates META-specific factors bind RNA in large abundance and are distinct from PCF and AMA-enriched RBPs. The relative quiescence of this stage as evidenced by Figs 1B and 5B, places extra emphasis upon the potential importance of META-enriched RNA binding proteins as promoters of transmission to mammalian hosts and differentiation to AMA forms [33]. Indeed, if the META stage cells cannot properly respond to environmental cues of a mammalian phagolysosome and differentiate to AMA stage, the lifecycle is halted and there is no infection or disease [2]. Of interest as well are the RNA binding factors distinguishing META stage parasites from AMA(MØ) RBPs as the former holds the potential to differentiate while the latter is in the midst of amastigogenesis (Fig 5B). Similarly, mRBPs which are enriched in AMA(MØ) relative to AMA(LD) cells may provide further candidate *trans-*regulators that control amastigogenesis as AMA(MØ) is still in the process while AMA(LD) have been through multiple replication cycles within this lifecycle stage long term.

Comparison of the predicted RBD-containing proteome of the *L.mexicana* genome to the isolated RBPome highlights some interesting distinctions including a decrease in the proportion of some zinc-finger domain proteins and in the basal translational machinery, particularly in the replicative stages. There are several, non-exclusive potential explanations for this. The temporally-defined isolation of these RBPomes may limit the scope. Although the RBPome has data from 4 distinct lifecycle stages, there are at least 3 lifecycle stages (nectomonad, leptomonad and haptomonad promastigotes) and multiple differentiation events not examined here in which non-isolated RBPs might be transiently expressed. Zinc finger proteins characteristically display reduced detection via mass spectroscopy. A study isolating and examining the missing lifecycle stages and differentiation time courses could detect additional RBPs, but isolation of these less-defined stages from sandflies at the numbers required for mass spectroscopy is experimentally prohibitive. Despite these caveats, this study presents the largest scale isolation of RBPs to date in kinetoplastids with valuable information on the types of proteins associating with mRNA in these parasites. These proteomes lend insight into the dynamic cellular context in which *Leishmania trans*-regulators bind RNA, as well as potential modifying enzymes which control RBP behavior and function.

Important findings from this work include the low correlation of protein expression versus transcript expression, the stage-specific variation in protein expression versus RNA binding potential, and the modulation of RNA binding protein enrichment during the *Leishmania* parasite lifecycle. This is the first study to examine the whole cell or mRNA binding proteome of the metacyclic promastigote parasites essential for human transmission, infectivity and lifecycle progression. We endogenously tagged and molecularly verified the association of multiple RBP candidates with distinct, stage-regulated transcript target pools. Functional investigations into candidate regulators will undoubtedly isolate factors essential for parasite lifecycle progression, viability and transmission. As the majority of kinetoplastid proteins are not homologous to other systems, including mammals, *trans*-regulators that enable the parasite to adapt and survive in different environments may provide viable targets for anti-leishmanial treatments.

## Experimental Procedures

### Parasite culture and purification

Early passage (P2-P3) PCF forms of *L. mexicana* strain M379 (MNYC/BZ/62/M379) were isolated from log phase (3-6e^6^ cells/mL) in M199 medium (pH 7.2) at 26°C [34]. META-enriched cultures were generated as previously described [20]. Briefly, PCF forms innoculated Grace’s media (10% hiFCS, 1% Pen/Strep, 1X BME vitamins, pH 5.5) at 1.5×10^5^ cells/ml and cultured for 7 days at 26°C. For J774.2 macrophage (MØ; (Sigma)) infection assays, META-enriched cultures were isolated via resistance to complement-mediated lysis in 20% human serum (Sigma) for 30min at 37°C and validated via *sherp* expression [21,23].

Validated META parasites were centrifuged and resuspended in complete DMEM medium (10% hiFCS, 1% Penicillin/Streptomycin, 2mM L-glutamine) and used to infect J774.2 MØ cultures at a 15:1 parasite/MØ ratio (1.25×10^7^ MØ/plate) for 6h at 34°C prior to washing 3X with prewarmed PBS, incubation in DMEM medium at 34°C for 18h, UV irradiation (or not), disruption of MØs and isolation of 24h post-infection (pi) intracellular amastigote forms (AMA(MØ)) using either a 45% Percoll or sucrose gradient as described [6].

### Mice infections

To isolate AMA(LD) parasites *in vivo*, 1×10^7^ serum-treated META *L.mexicana* cells were injected in rumps of 8-12 week old Balb/c mice (Charles River, UK). After 4 months the mice were sacrificed and the lesions harvested as described [6,24]. For ethical reasons, AMA(LD) were harvested only for XL mRNP samples.

### Isolation of *Leishmania* mRNA-bound proteomes

For the mRNA interactome capture experiments, all conditions were equivalent for the *Leishmania* lifecycle forms PCF, META, AMA(MØ) and AMA(LD) between XL and nonXL samples except the irradiation. *In vivo* UV-crosslinking was performed using the LT40 “Minitron” system (UV03 Ltd.) [28]. Cells at 5×10^6^ cells/ml confluence (~0.6 OD) were irradiated for 120s (Fig 2B; ~1.6 mJ/cm^2^) for optimal *in vivo* RBP:RNA (mRNP) crosslinking with superior mRNA integrity and negligible heat stress compared to a standard Stratalinker which exposes the cells to 100X the heat (~150 mJ/cm^2^) [28]. Both AMA(MØ) and AMA (LD) were irradiated in host MØs, which were then homogenized to release parasites for gradient purification as above.

Triplicate samples of ~5×10^9^ cells of each lifecycle stage (XL and nonXL) were resuspended in 15ml Lysis buffer [35] (20mM Tris-HCl, pH7.5, 500 mM LiCl, 0.5% LDS, 1mM EDTA and 5mM DTT with cOmplete EDTA-free Protease Inhibitors ™ (Roche)) for 10min at 0°C, passed through a 25G needle until clear, centrifuged 10min at 4000*g* and incubated with oligo(dT)_25_ beads 30min at 4°C (New England Biolabs). Remaining steps for polyA RNA isolation were performed as described [18]. mRNA-bound proteins were precipitated via TCA precipitation, and the resulting protein concentration measured using Micro BCA Protein assay kit (Thermo).

### Label-free Quantitative Mass Spectrometry Analysis

#### Trypsin Digestion

Triplicate biological samples were solubilised in NuPAGE LDS sample buffer (Life Technologies), heated at 70°C for 10min and ran on a 7cm NuPAGE Novex 10% Bis-Tris gel (Life Technologies) at 200 V for 6min. Gels were stained with SafeBLUE protein stain (NBS biologicals) for 1hr before destaining with ultrapure water for 1hr. In-gel tryptic digestion was performed after reduction with dithioerythritol and S-carbamidomethylation with iodoacetamide. Gel pieces were washed two times with aqueous 50% (v:v) acetonitrile containing 25mM ammonium bicarbonate, then once with acetonitrile and concentrated in a vacuum for 20min. Sequence-grade, modified porcine trypsin (Promega) was dissolved in 50mM acetic acid and diluted with 25mM ammonium bicarbonate to give a final trypsin concentration of 0.02g/L. Gel pieces were rehydrated with 25L of trypsin solution, incubated for 10 min then 25mM ammonium bicarbonate solution was added to cover the gel pieces. Digests were incubated overnight at 37°C. Peptides were extracted by washing three times with aqueous 50% (v:v) acetonitrile containing 0.1% (v:v) trifluoroacetic acid, before concentrating in a vacuum and reconstituting in aqueous 0.1% (v:v) trifluoroacetic acid. A common sample pool was created by taking equal aliquots of all samples.

#### LC-MS/MS

Samples were loaded onto an UltiMate 3000 RSLCnano HPLC system (Thermo) equipped with a PepMap 100Å C18, 5µm trap column (300µm x 5mm Thermo) and a PepMap, 2µm, 100Å, C18 EasyNano nanocapillary column (75m x 150mm; Thermo). The trap wash solvent was aqueous 0.05% (v:v) trifluoroacetic acid; trapping flow rate was 15µL/min. The trap was washed for 3min before switching flow to the capillary column. Separation used gradient elution of two solvents: solvent A, aqueous 1% (v:v) formic acid; solvent B, aqueous 80% (v:v) acetonitrile containing 1% (v:v) formic acid. The flow rate for the capillary column was 300nL/min and the column temperature was 40°C. The linear multi-step gradient profile was: 3-10% B over 7min, 10-35% B over 30min, 35-99% B over 5min and then proceeded to wash with 99% solvent B for 4min. The column was returned to initial conditions and re-equilibrated for 15min before subsequent injections. The nanoLC system was interfaced with an Orbitrap Fusion™ Hybrid™ mass spectrometer (Thermo) with an EasyNano ionisation source (Thermo). Positive ESI-MS and MS2 spectra were acquired using Xcalibur software (version 4.0, Thermo). Instrument source settings were: ion spray voltage, 1,900V; sweep gas, 0 Arb; ion transfer tube temperature; 275°C. MS 1 spectra were acquired in the Orbitrap Fusion™ with: 120,000 resolution, scan range: *m/z* 375-1,500; AGC target, 4e^5^; max fill time, 100ms. Data dependant acquisition was performed in top speed mode using a 1s cycle, selecting the most intense precursors with charge states. Easy-IC was used for internal calibration. Dynamic exclusion was performed for 50s post precursor selection and a minimum threshold for fragmentation was set at 5e^3^. MS2 spectra were acquired in the linear ion trap with: scan rate, turbo; quadrupole isolation, 1.6 *m/z*; activation type, HCD; activation energy: 32%; AGC target, 5e^3^; first mass, 110 *m/z*; max fill time, 100ms. Acquisitions were arranged by Xcalibur to inject ions for all available parallelizable time.

### Fusion PCR and endogenous 3xHA tagging of N-termini

For the generation of 3xHA N-tagged cell lines, the original modular pPOTv2 vector (Dyer et al. 2016) was modified. The Ty-GFP-Ty tag was excised (HindIII*/*BamHI) and replaced by a 3xHA tag. Fusion PCR was performed as previously described. For transfections, 2×10^7^ PCF cells were resuspended in Tb-BSF buffer (100µl) and transfected [36]. The parasites were selected with 10µg/ml Blasticidin (Sigma).

### RNA co-immunoprecipitation and qRTPCR

HA-tagged RBPs were immunoprecipitated from PCF or META lysates with anti-HA magnetic beads (Thermo) co-immunoprecipitated RNA were extracted and measured via qRTPCR using SuperScript IV Reverse Transcriptase, Fast SYBR Green Master Mix and Quantstudio 3 PCR System (Thermo Fisher) as previously described [35]. Relative levels of qRTPCRs were calculated via 2^−ΔΔCt^ using *nmt* as a constitutive control [27].

For the mRNA-bound capture experiment, the quality of the results was measured using depleted (*18S* ribosomal gene) and non-depleted (*nmt*) relative mRNA values before and after oligo(dT)_25_-labeled magnetic bead purification. The relative levels were measured using the 2^−ΔCt^ method, using total RNA from samples prior to mRNA purification as a reference. The full list of primers used is provided in Supplementary S3 Table.

### Western blot and protein expression

The different lifecycle stages of 3xHA endogenously tagged cell lines were grown and purified as described above. Parasites were lysed in Laemmli buffer and samples separated by SDS-PAGE, transferred to PVDF, labeled with anti-HA (1:10,000, Pierce) and anti-mouse IgG-HRP (1:50,000, Sigma) and developed using ECL (GE Healthcare). Relative protein expression was quantified using ImageJ software.

### Data Analysis

Peak lists in .raw format were imported into Progenesis QI and LC-MS runs aligned to the common sample pool. Precursor ion intensities were normalised against total intensity for each acquisition. A combined peak list was exported in .mgf format for database searching against *L.mexicana* sequences appended with common proteomic contaminants (8,365 sequences). MascotDaemon (version 2.5.1, Matrix Science) was used to submit the search to a locally-run copy of the Mascot program (Matrix Science Ltd., version 2.5.1). Search criteria specified: Enzyme, trypsin; Fixed modifications, Carbamidomethyl (C); Variable modifications, Oxidation (M); Peptide tolerance, 5ppm; MS/MS tolerance, 0.5Da; Instrument, ESI-TRAP. Search results were filtered to require a minimum expect score of 0.05. The Mascot .XML result file was imported into Progenesis QI and peptide identifications associated with precursor peak areas. Relative protein quantification was derived from unique peptide precursor ion intensities. Accepted quantifications were required to contain a minimum of two unique peptides. Statistical testing was performed in Progenesis QI and ANOVA-derived p-values were converted to multiple test-corrected q-values using the Hochberg and Benjamini approach. Final quantification results were stripped of non-*Leishmania* spp. identifications for brevity.

### Bioinformatics analysis

#### Comparison of predicted versus isolated RBPomes

A list of characterized, published RNA binding domains was compiled from the InterPro database using manual curation and the search term ‘RNA binding’. The 1,407 isolated RBP candidates (1,638 LC MS/MS protein identities) of the *L.mexicana* XL-RBPome were compared with the compiled list on Tritrypdb.org (08/2018) [37] using the ‘InterPro domain’ function with the condition ‘intersect’ (New search/Protein features and properties/InterPro domain). This same method of comparison was also completed between the compiled list and the whole *L.mexicana* protein coding genome. Diagrams were created with Microsoft Excel 2013, Corel Draw 2017 and Inkscape 0.92.3 softwares.

#### Venn diagrams

WC and XL proteomes at each stage were filtered to include proteins with mean intensity values across each of the 3 replicates per condition intensity ≥ 10^6^. These proteins were used to create Venn diagrams in Python 2.7 using Matplotlib v.1.5.3 and Matplotlib-Venn package version 0.11.5.

#### Gene Ontology term analysis

Molecular Function GO Terms significantly enriched in the isolated mRBPome relative to the predicted *L.mexicana* proteome were derived using Tritrypdb.org (04.2017) [37]. REVIGO software was used to refine and visualize enriched terms (revigo.irb.hr) [38].

#### Volcano plots

Volcano plots were generated in Python 2.7 using Matplotlib v.1.5.3 and mpld3.js v.1.0 for interactive visualization. Log 2 fold-change values for every protein were calculated by taking the log 2 value of the ratio of mean intensity value + 1 between conditions. Independent t-tests were conducted using intensity values in each of the 3 replicates per condition to generate the p-values used in the volcano plots.

#### Whole cell proteome versus transcriptome [25] data correlation

WC proteome and transcriptome correlations were made by averaging intensity values across replicates, using genes of q value ≤ 0.05 with at least 2 peptide hits. This was done for both the WC proteome versus the whole transcriptome, to identify any overarching correlations, and for each specific stage. For WC proteome and transcriptome comparisons, the ratio of PCF to AMA(MØ) intensity was calculated for each replicate, averaged and then log_2_ transformed. Genes obtained from the WC proteome were then compared to transcripts previously found to be differentially expressed between AMA(MØ) and promastigote^log^ stages [25]. Intersecting genes between both datasets were then plotted against each other, using the ggplot2 R package, and the regressions fitted in Fig 5A were modeled based on a loess linear model that derive the statistics in Table 1 below.

**Table 1:** RNA-bound versus Whole Cell proteome statistics and parameters. Statistics associated with WC and mRNA-associated (XL) Proteome comparisons of PCF, META and AMA lifecycle stage *L.mexicana* parasites (Fig 5B).

| Lifecycle Stage | UV crosslinked proteome vs whole cell proteome<br>(simple linear model based on XL vs WC correlation) |  |  |
| --- | --- | --- | --- |
|  | Standard Error (d.f.) | Adjusted R2 | p value |
| All stages | 3.959 (4,219) | 0.0066 | 7.05E-08 |
| PCF | 2.956 (1,405) | 0.0118 | 2.64E-05 |
| META | 4.051 (1,405) | 0.1325 | < 2.2E-16 |
| AMA | 3.486 (1,405) | 0.0178 | 3.11E-07 |

Following from this, the expression values from the proteomics data were then compared for each life cycle stage in both XL and WC conditions. As before, the intensity values were averaged between replicates, and were plotted using ggplot2, however these were instead fit with a simple linear regression model, as a loess model did not improve the fit to the data.

### Experimental Design and Statistical Rationale

Biological triplicates of each proteome sample (WC, XL and nonXL) at all stages (PCF, META, AMA (Mø) and AMA(LD)) were quantitatively assessed for peptide ion enrichment. Data were searched against the TriTrypDB *Leishmania mexicana* proteome (version 8.1 – 30^th^ Sep 2014, 8,250 sequences; 5,180,224 residues) concatenated with 115 common proteomic contaminants including trypsin and human keratins (38,188 residues). Enzyme search parameters required full trypsin specificity (cleavage C-terminal to K or R not preceding P) and allowed for up to one missed cleavage. An identification false discovery rate of 2.4% was estimated empirically by searching against a reverse database and comparing the relative proportion of matches, restricting to the top scoring identification for each MS2 query and requiring a minimum of two unique peptides for each accepted protein.

## Acknowledgements

This work was supported by the Medical Research Council [grant number MR/L00092X/1 to PBW]. Funding for open access charge: Medical Research Council and University of York. LCMS within the York Centre of Excellence in Mass Spectrometry was founded through Science City York and Yorkshire Forward/Northern Way Initiative; and is supported by EPSRC [grant numbers EP/K039660/1, EP/M028127/1]. All of the Proteomic work and part of the Bioinformatic analyses was conducted within the Bioscience Technology Facility (https://www.york.ac.uk/biology/technology-facility) at the University of York. UV-crosslinking was advised by the Tollervey and Granneman labs. We would like to thank Deborah Smith, Alvaro Acosta-Serrano and Dawn Coverley for helpful comments on this manuscript.

## Ethics statement

All animal procedures for this research have been passed by the University Animal Welfare Ethical Review Body (AWERB) and is carried out under the Home Office Project Licence [39_4377]. The University of York and the Home Office Project Licence meets all the standard conditions and works according to Codes of Practice under the under the Animals (Scientific Procedures) Act 1986 (Amended 2012).

## Data availability

Proteomic data sets are available to download from MassIVE (MSV000083023) and ProteomeXchange (PXD011340). All proteomic data has been submitted to TriTrypDB (http://tritrypdb.org/), which is possible through the collaborative efforts between EuPathDB, GeneDB and the Center for Infectious Disease Research (CIDR).

## Supporting information captions

**Figure S1.**
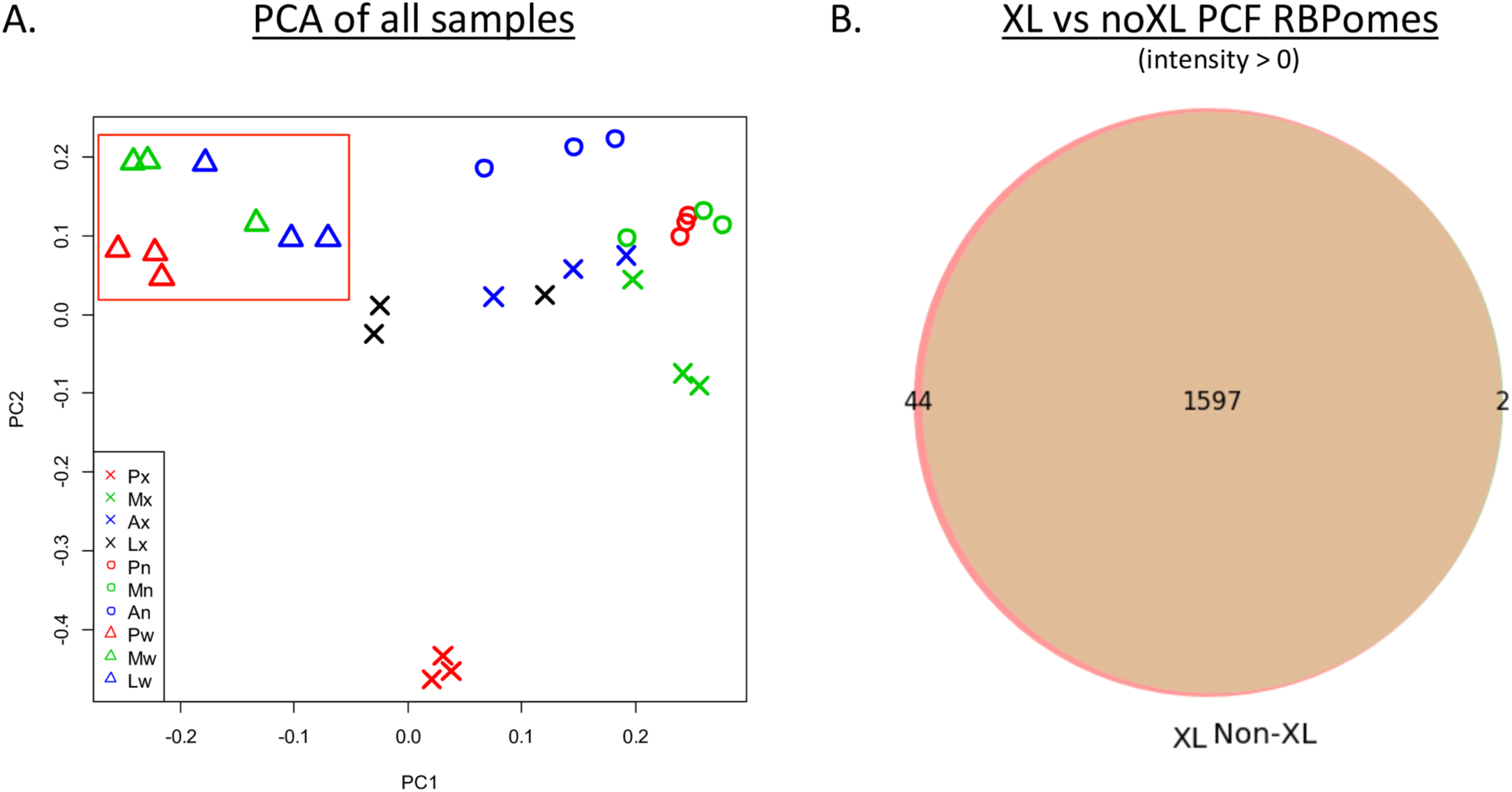
XL and nonXL RBPomes are distinct from whole proteomes and nearly identical. A. PCA analyses of all Proteome data. Whole proteomes segregate from RBPomes regardless of lifecycle stage. B. Venn Diagram of XL vs nonXL RBPomes without filtering for differential enrichment. Close overlap reveals isolated XL and nonXL RBPome protein identities are nearly identical.

**Figure S2.**
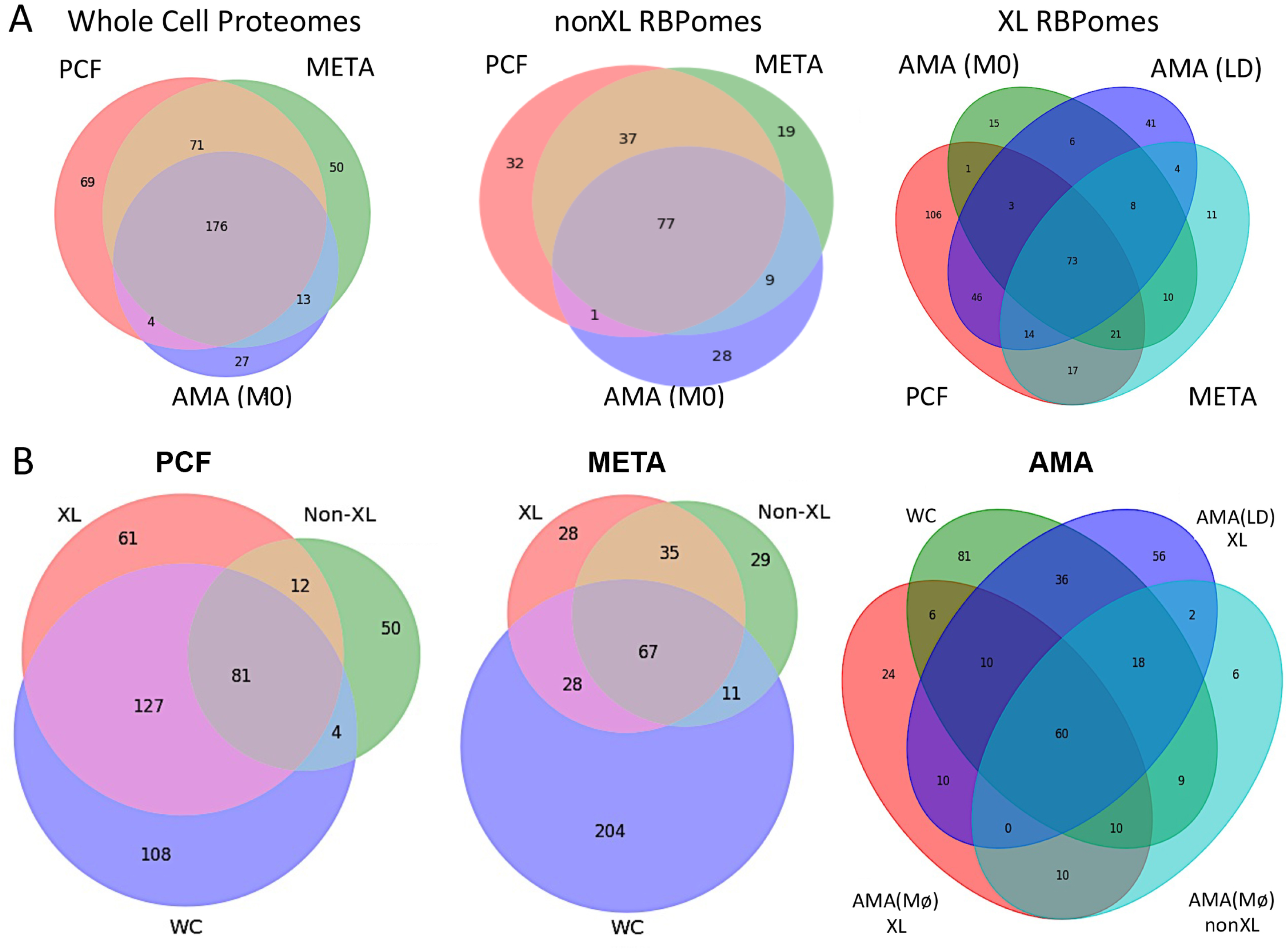
Venn Diagram comparison of enriched proteomes across the *Leishmania* lifecycle. (A) Protein enrichment analyses compare each lifecycle stage of the WC, XL and nonXL proteomes. Proteomes of each lifecycle stage were filtered to select highly enriched proteins with mean intensity values across each of the 3 replicates ≥ 10^6^. These enrichment-selected proteins were used to create Venn diagrams in Python 2.7 using Matplotlib v.1.5.3 and Matplotlib-Venn package version 0.11.5. (B) Complementary Venn diagrams comparing the WC, XL and nonXL proteomes of each lifecycle stage.

**Figure S3.**
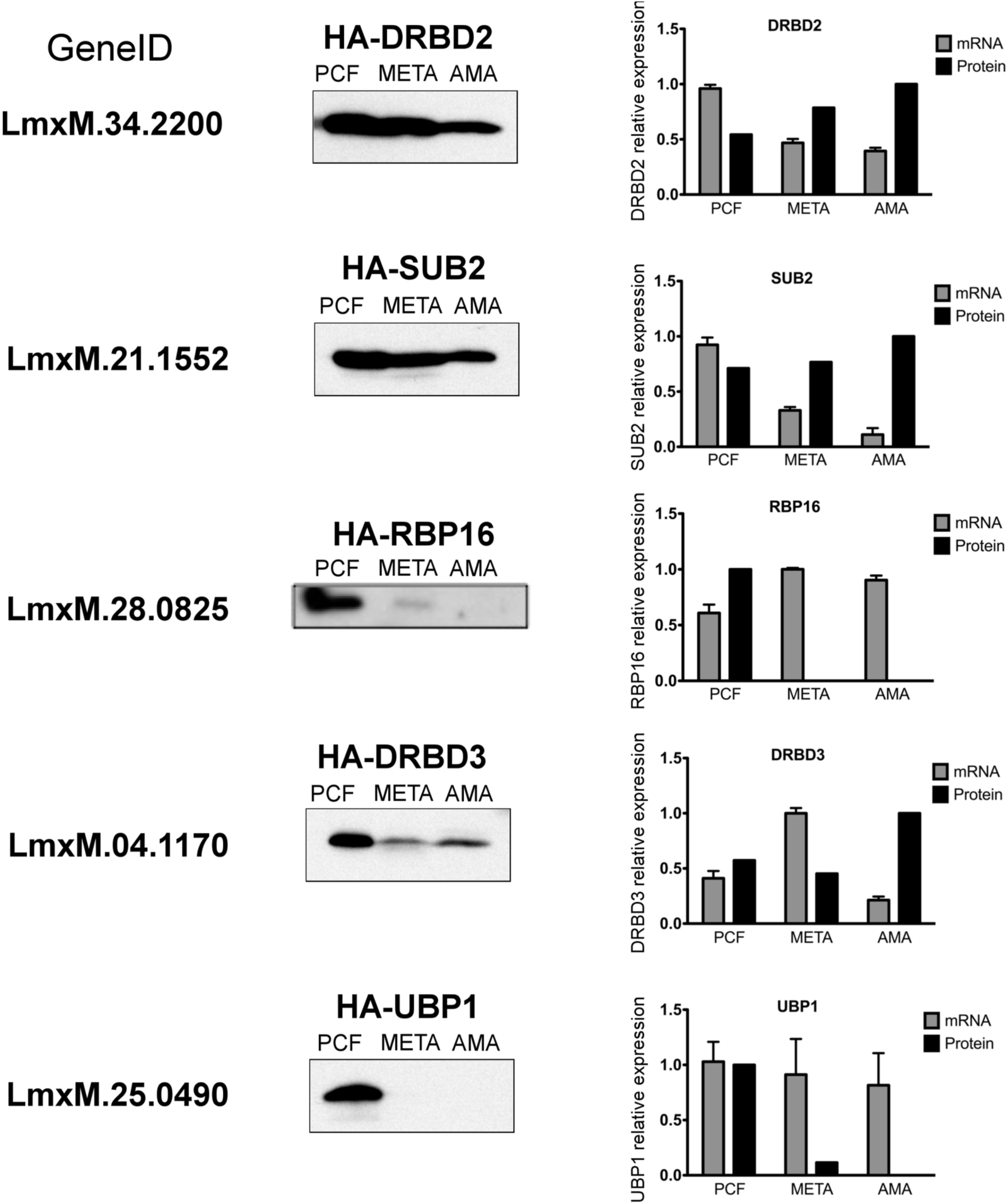
Relative RBP protein and mRNA expression through lifecycle progression. Relative expression of endogenously-tagged LmxM.34.2200 (DRBD2), LmxM.21.1552 (SUB2), LmxM.28.0825 (RBP16), LmxM.04.1170 (DRBD3) and LmxM.25.0490 (UBP1) is shown via anti-HA western blot and qRTPCR analyses. Levels of each were quantified relative to NMT protein and transcript expression [26].

**Table S1. Relative intensity of filtered proteomes.** Accession numbers and relative intensities (percentage) of RBPome proteins isolated in XL, nonXL and WC proteomes filtered for triplicate-consistent, high quality reads with at least 2 unique peptides are provided. Red to Blue color gradient is indicative of relative intensity values. Intensity sum for each row = 100%. Relative Percent Intensity values were calculated by expressing normalized intensity values for each protein as the percentage of the sum of normalized intensity values among all samples. Q value represents significance of intensity shift between different conditions of each triplicate. Presented q-values are Hochberg and Benjamini multiple-test corrected false discovery rates calculated from Progenesis QI ANOVA tests for significant difference between sample groupings. Q-values <0.05 are highlighted in green. Identification descriptions are approximate and predominantly homology-based as the overwhelming majority of factors are uncharacterized in this system.

**Table S2. Label-free Quantification Summary.** Table of Progenesis QI-derived label-free quantified proteins detailing relative raw protein abundances observed among all samples. Columns headed Normalized Intensity list summed MS^1^ peak areas extracted from non-conflicting peptides post normalisation against total ion intensity. Statistical analyses are as in Table S1.

Table S3. Primers used in experimental procedures.

